## Supplementary Note, Figures, and Tables for "Differential regulation of alternative promoters emerges from unified kinetics of enhancer-promoter interaction"

### 1. 3’RACE and PCR

3’RACE and PCR were used to verify the difference in 3’-end sequences between P1- and P2-driven transcripts (Supplementary Fig. 1c). Total RNA was extracted from 2-hour-old embryos (collected at 25°C) using PureLink RNA Mini Kit (12183020, Invitrogen) with on-column DNase treatment to remove genomic DNA. The 1$^{st}$ strand cDNA was reverse-transcribed using PrimeScript Double Strand cDNA Synthesis Kit (6111A, Takara) with an Oligo(dT) primer (Supplementary Table 2). After destroying the mRNA template with RNase H (AM2292, Invitrogen), the 2$^{st}$ strand cDNA was synthesized using DNA Polymerase (12369010, Invitrogen) with P1- and P2-specific RT primers (designed to cross the exon-exon junction, Supplementary Table 2), respectively. By removing the remaining single-stranded cDNAs with S1 Nuclease (18001016, Invitrogen, the reaction can be stopped by adding 2 μL of 0.5 M EDTA and heating at 70 °C for 10 minutes), a 34-cycle PCR was performed with promoter-specific forward primers and 3’end-specific backward primers (designed to be upstream and downstream of *hb*-RB 3’end, respectively) to amplify P1- and P2-specific cDNAs (Supplementary Table 2). To verify that no genomic DNA existed in the RT products, we also used a pair of PCR primers targeting the non-transcribed sequences upstream of the *hb* gene. PCR products were loaded onto 1% agarose gel for electrophoresis. The gel results confirmed that P2-driven transcripts terminate at a more distant site than P1-driven transcripts (Supplementary Fig. 1c).

### 2. HCR-FISH probe design and labeling

HCR-FISH labels the target transcripts with two sets of probes[1,2]: (1) a set of primary probes, each containing a 20-nt sequence complementary to the target transcripts, a 5-nt spacer, and a 36-nt HCR initiator on the 3’ end; (2) a set of two fluorophore-conjugated amplifier probes that labels the initiator with HCR amplification. In this study, pre-designed amplifier probe sets, B3 and B5[2] (Supplementary Table 3), were synthesized (Thermo Fisher Scientific) with fluorophore tags (Alexa Fluor™ 488 for B3 probes and Alexa Fluor™ 594 for B5 probes). Primary probe sets were designed and synthesized (Sangon Biotech) for *yellow* and *lacZ* transcripts (nine probes with B3 initiator for *yellow* and ten probes with B5 initiator for *lacZ*, Supplementary Table 3).

smFISH and HCR-FISH co-staining of embryos was performed according to a previously published protocol[1,2]. Briefly, fixed embryos were rehydrated (4 × 10 min) in PBST (1× PBS, 0.1% (v/v) Tween-20) at room temperature, washed (1 × 10 min) in hybridization buffer (5× SSC, 30% (w/v) formamide, 9 mM citric acid pH 6, 50 μg/mL heparin, 1× Denhardt’s solution, 10% dextran sulfate, and 0.1% (v/v) Tween-20) at room temperature, and incubated with smFISH probes and HCR-FISH primary probes (in hybridization buffer) at 37°C overnight. After hybridization, embryos were washed three times (15 min each) at 37°C in 75%, 50%, and 25% probe wash buffer (5× SSC, 30% (w/v) formamide, 9 mM citric acid pH 6, 50 μg/mL heparin, and 0.1% (v/v) Tween-20) diluted in 5× SSCT (5× SSC, 0.1% (v/v) Tween-20), respectively, and further washed twice (15 min each) in 5× SSCT at room temperature. For HCR amplification, each amplifier probe was separately heated to 95 °C for 90 seconds and cooled to room temperature for 30 min to open the hairpin structure. Embryos were washed in amplification buffer (5x SSC, 0.1% Tween-20, 10% dextran sulfate) at room temperature for 30 min and incubated with the amplifier probe set (in amplification buffer) at room temperature for 4 hours. Following HCR amplification, embryos were further washed (4 × 10 min) in 5x SSCT at room temperature and counterstained with Hoechst 33342 for 10 min at room temperature. Following additional washes (4 × 10 min) in 5x SSCT, embryos were mounted in Aqua-Poly/Mount (Polysciences, 18606). Imaging was performed after the samples were completely solidified.

**3. Quantifying the relationship between different probe signals in promoter deletion embryos**

To determine the relationship between the nascent signals of the CDS and promoter-specific probes for each promoter, we co-labeled promoter mutant embryos (ΔP1C and ΔP2C)[3] with P1-5'UTR, P2-3'UTR, and CDS probes (as well as a *lacZ* probe set for identifying homozygous embryos, see Methods). For each embryo, we computed the nuclear nascent signal of each probe set (in units of the number of mRNA molecules) and binned individual data points by nuclear position (Supplementary Fig. 4a). We observed no P1-5'UTR signal for ΔP1C and no P2-3'UTR signal for ΔP2C, consistent with the embryos' genotypes. The average nascent signals of 5' probes (CDS for ΔP1C, P1-5'UTR for ΔP2C) were much larger than that of 3' probes (P2-3'UTR for ΔP1C, CDS for ΔP2C), agreeing with theory (see Supplementary Note 6.5). By plotting the binned nascent CDS signal against promoter-specific probe signals (0.3–0.6 EL), we showed that the two were roughly in proportion (Supplementary Fig. 4b), confirming a theoretical result (see Supplementary Note 6.5). Specifically, a linear regression estimated a ratio of ~5.6 between CDS and P2-3'UTR for ΔP1C, and a ratio of ~0.4 between CDS and P1-5'UTR for ΔP2C. These values are roughly similar to $a_2$ and $a_1$ estimated from the WT embryo. Based on these ratios, the post-elongation residence time ($T_R$) was estimated to be $T_{R\text{-}P2}$ = 10 s for ΔP1C and $T_{R\text{-}P1}$ = 47 s for ΔP2C. Both values were smaller than those in the WT embryo, suggesting that transcripts from the two promoters may interrupt each other's termination process. The remaining difference between $T_{R\text{-}P1}$ and $T_{R\text{-}P2}$ in single-promoter embryos may be intrinsic since P1- and P2-driven transcripts have different termination sites.

Moreover, with colocalization analysis, we showed that 5' probes (CDS for ΔP1C, P1-5'UTR for ΔP2C) labeled more nascent mRNA foci than their 3' counterparts (P2-3'UTR for ΔP1C, CDS for ΔP2C) (Supplementary Fig. 4c). This result indicates that most CDS-only foci in the WT embryo are P2-driven. It explains the difference between the CDS and P2-3'UTR signals observed in the paper (Fig. 1c–f).

### 4. Determining the accuracy of Bcd quantification using immunofluorescence

Quantifying Bcd gradient in the embryo is critical to understanding the mechanism of *hb* regulation. Such quantification typically relies on live imaging of transgenic embryos with Bcd-EGFP or immunofluorescence imaging of fixed embryos[4,5]. However, it is always a concern whether the measured gradient represents the functional Bcd signal. Specifically, the decay length of the Bcd-EGFP gradient measured from live imaging (~20% EL) differs significantly from that of the functional gradient (15.0% ± 1.4% EL) estimated from the shift of gene expression boundary in response to Bcd dosage change[6]. According to previous studies, this difference is mainly due to the slow maturation of EGFP[7,8]. Since Bcd molecules are synthesized at the anterior pole of the embryo, many anterior Bcd-EGFPs may be too young to be visible. As a result, the observed Bcd gradient is milder than the actual one. This effect was estimated to cause ~15% increase in the measured decay length[7]. Thus, quantitative application of live Bcd data typically requires a correction of the EGFP maturation effect[7,9,10].

On the other hand, immunofluorescence measurements do not require EGFP and thus typically reported significantly smaller length constants (~16% EL[7,11,12], with few exceptions[5]). In this work, we used immunofluorescence to label Bcd in fixed WT embryos. The estimated decay length is 15.9% ± 0.5% EL (Supplementary Fig. 6b), which agrees well with the functional decay length estimated in a recent work[6]. It suggests that our immunofluorescence data reflects the functional Bcd gradient.

**5. Speculating Bcd binding dynamics using immunofluorescence**

The binding of transcription factors on the regulatory sequence is the critical step of transcriptional regulation. Previous live imaging studies of GFP fusion proteins revealed that Bcd and other transcription factors combinatorically bind DNA to form dynamic clusters, which transiently interact with Bcd-target genes to activate transcription[13,14]. However, accurate quantification of Bcd binding dynamics using live imaging is still technically challenging to date due to the low signal level and slow maturation of the fluorescence protein. In this work, we measured local enrichment of Bcd at P1- and P2-active loci in fixed embryos, which corresponds to a snapshot of Bcd binding in the vicinity of *hb* loci at the time of fixation. Although our measurement did not directly capture the temporal information of Bcd binding, it is accurate enough for estimating the number of bound Bcd molecules. By computing the average level of Bcd binding in different Bcd concentration ranges, we identified two typical Bcd binding states for *hb* activation. By relating these binding states with the statistics of promoter transcription, we reconstructed the dynamics of Bcd binding and *hb* promoter activation using a mathematical model. Such understanding of Bcd binding dynamics may inspire future research of transcription factor binding.

### 6. Mathematical modeling of transcriptional kinetics

#### *6.1 Model selection*

Transcription kinetics determines the nascent mRNA copy number distribution on individual gene loci. Specifically, bursty gene expression often results in a multimodal distribution[15,16]. A widely used model for describing bursty gene expression is the two-state telegraph model[15,17-20], in which the gene randomly switches between an inactive and an active transcription state. If transitions between states are slower than the residence time of nascent mRNA on the transcription site, the nascent mRNA distribution predicted from the model exhibits two Poissonian peaks[20]. Generalizing the model to include more states can create more peaks in the distribution[21]. As a general feature, the number of peaks (i.e., modality) of the experimentally observed distribution sets the minimum number of gene states required for modeling the transcription process (see Supplementary Note 6.3 for details).

In this study, the distributions of P1 and P2 nascent mRNA signals both exhibited trimodal distributions (Fig. 5a). One peak in the distribution corresponds to silent loci ($m$ = 0), while two other peaks correspond to two groups of active loci with different expression levels. There are two possible explanations for this phenomenon: (1) individual promoters perform three-state transcription kinetics, and (2) each observed promoter locus is composed of a pair of closely located sister loci indistinguishable under the microscope[22,23]. To evaluate these two explanations, we plotted the distribution of nascent mRNA signals measured from individual optically resolved sister loci (Fig. 5b). For each promoter, the sister loci exhibited two active populations. The distribution was well fitted by a sum of two Poisson distributions (Considering the intensity threshold used for identifying active transcription sites, the very left part of the distribution (<3 mRNAs) was neglected). By comparing the weights of the two Poisson peaks, we showed that the minor population in the distribution corresponds to 38.4% ± 3.3% of P1 and 17.0% ± 6.1% of P2 sister loci (mean ± s.e.m., data from seven embryos at nc13). Thus, the activity of a single promoter needs to be described with at least three transcription states. Moreover, since the trimodal distributions of P1 and P2 nascent mRNAs each have a peak at $m$ = 0 and two distinct peaks at $m > 0$, we speculated that one of the three transcription states is inactive, while two others are active with distinct rates of transcription.

Our model is different from the three-state models proposed in some other papers[24,25], which contained two inactive states and one active state. Those "2 OFF/1 ON" models can describe fine steps within the inactive phase of gene regulation. However, their nascent mRNA distributions were either unimodal or bimodal. I.e., they cannot reproduce the trimodal distribution observed in this paper (see Supplementary Note 6.3). In fact, when fitting the experimental distributions, we used a more general model, in which $k_{\mathrm{INI},1}$ and $k_{\mathrm{INI},2}$ were allowed to reach zero (see Supplementary Note 6.10). The inferred $k_{\mathrm{INI},1}$ and $k_{\mathrm{INI},2}$ were always nonzero, indicating that the "2 ON/1 OFF" model is the right choice for *hb* promoters.

It should be noted that both the "2 ON/1 OFF" and "2 OFF/1 ON" models are simplifications of the actual transcription process, which involves far more molecular steps. Since our paper focuses on the many-body interactions between enhancers and promoters, whose primary feature has been captured by two active states, it is valid to neglect detailed kinetic steps within the inactive state.

#### *6.2 Model assumptions*

The nascent transcription of each promoter locus was modeled as a three-state process. The model considers three transcription states of the promoter: an "OFF" state (denoted as state 0), where the promoter is transcriptionally inactive, and two "ON" states (denoted as states 1 and 2), where the promoter actively initiates new transcripts. State transitions and mRNA initiations are assumed to be Poisson processes with specific rates $k_{ij}$ and $k_{\mathrm{INI},i}$ ($i$, $j$ = 0, 1, 2), respectively. Following initiation, each nascent mRNA molecule elongates to the final length $L$ with a constant speed $V_{\mathrm{EL}}$. Once completed, the mRNA resides on the gene for an extra termination period, $T_{\mathrm{R}}$, before being released.

At a given observation time, the state of the system at a given observation time $t_{\mathrm{ob}}$ is determined by the promoter state $n$ ($n$ = 0, 1, 2) and the total signal of nascent mRNA $m$ ($m \geq 0$). Since a nascent mRNA molecule stays on the gene for a fixed period $T_{\mathrm{RES}} = L/V_{\mathrm{EL}} + T_{\mathrm{R}}$, $m$ is the sum of signals from all transcripts initiated between $t_{\mathrm{ob}} - T_{\mathrm{RES}}$ and $t_{\mathrm{ob}}$, i.e., $m = \sum_{-T_{\mathrm{RES}} \leq \tau \leq 0} g(\tau)$. Here, $\tau = t - t_{\mathrm{ob}}$ is the time relative to $t_{\mathrm{ob}}$. Considering that nascent transcripts may be incomplete, we defined a contribution function $g(\tau)$ to describe the signal from a transcript initiated at time $\tau$ [19,20]. $g(\tau)$ varies between zero and one, and its exact shape depends on the target positions of the probe set and the magnitude of $T_{\mathrm{R}}$. Specifically, $g(\tau) = 1$ for $-T_{\mathrm{RES}} \leqslant \tau \leqslant -L/V_{\mathrm{EL}}$, while it decreases gradually from one to zero for $-L/V_{\mathrm{EL}} < \tau \leqslant 0$. The shape of $g(\tau)$ for each probe set is shown in Supplementary Fig. 5b.

#### *6.3 Master equation*

For a general $N$-state model, the master equation for the probability distribution of ($n$, $m$) is

$$\frac{d\mathbf{P}(m)}{d\tau} = (\mathbf{K} - \mathbf{K}_{\mathrm{INI}})\mathbf{P}(m) + \mathbf{K}_{\mathrm{INI}}\mathbf{P}(m - g(\tau)) \quad (1)$$

where $\mathbf{P}(m)=\begin{bmatrix} P(0,m) \\ P(1,m) \\ \vdots \\ P(N-1,m) \end{bmatrix}$ denotes the nascent mRNA distributions of each transcription state,

$\mathbf{K}=\begin{bmatrix} -\sum_{i\neq 0} k_{0,i} & k_{10} & \cdots & k_{N-1,0} \\ k_{01} & -\sum_{i\neq 1} k_{1,i} & \cdots & k_{N-1,1} \\ \vdots & \vdots & \ddots & \vdots \\ k_{0,N-1} & k_{1,N-1} & \cdots & -\sum_{i\neq N-1} k_{1,i} \end{bmatrix}$ describes transitions between different promoter states,

and $\mathbf{K}_{\mathrm{INI}}=\begin{bmatrix} k_{\mathrm{INI},0} & 0 & \cdots & 0 \\ 0 & k_{\mathrm{INI},1} & \cdots & 0 \\ \vdots & \vdots & \ddots & \vdots \\ 0 & 0 & \cdots & k_{\mathrm{INI},N-1} \end{bmatrix}$ describes transcription initiation of each promoter state[20,26].

The solution of the equation varies with the magnitudes of $k_{i,j}$ and $k_{\mathrm{INI},i}$. Specifically, in an extreme case of slow state-transitions (i.e., $\mathbf{K}T_{\mathrm{RES}} \approx 0$), each row in Equation (1) is decoupled into a 1-state equation. The steady-state solution of each 1-state equation is a unimodal distribution peaked at $m_{\mathrm{peak},i} \propto k_{\mathrm{INI},i}$[15,20,26]. Thus, the steady-state solution of Equation (1) is a linear combination of these unimodal distributions weighted by the probability of each transcription state. Obviously, for a system with $N$ distinct $k_{\mathrm{INI},i}$, the solution is a multimodal distribution with $N$ peaks, each of which corresponds to a transcription state. If $k_{\mathrm{INI},i}$ of some states are identical, the number of peaks reduces accordingly. Moreover, outside the slow-transition region of the parameter space, coupling between rows in Equation (1) tends to merge peaks and further reduces the distribution modality[20,21,26]. Consequently, the steady-state solution of an $N$-state model with $M$ distinct transcription initiation rates has, at most, $M$ peaks. This result may be used as a criterion for model selection (see Supplementary Note 6.1 for details).

In this study, we applied Equation (1) to the three-state system ($N$ = 3) with $\mathbf{P}(m)=\begin{bmatrix}P(0,m)\\P(1,m)\\P(2,m)\end{bmatrix}$,

$\mathbf{K}=\begin{bmatrix}-k_{01}-k_{02} & k_{10} & k_{20}\\ k_{01} & -k_{10}-k_{12} & k_{21}\\ k_{02} & k_{12} & -k_{20}-k_{21}\end{bmatrix}$, and $\mathbf{K}_{\mathrm{INI}}=\begin{bmatrix}0 & 0 & 0\\ 0 & k_{\mathrm{INI},1} & 0\\ 0 & 0 & k_{\mathrm{INI},2}\end{bmatrix}$. Here, state 0 is set to be inactive ($k_{\mathrm{INI},0}=0$), while states 1 and 2 are allowed to be active ($k_{\mathrm{INI},1} \neq k_{\mathrm{INI},2} \geqslant 0$). Assuming that the marginal distribution of the promoter state at $\tau = -T_{\mathrm{RES}}$ is $q(n)$, we can apply an initial condition of $\mathbf{P}(m) = q(n)\delta_{m,0}$ to solve Equation (1) for $\mathbf{P}(m)$ at $\tau = 0$. Specifically, $q(n)$ at steady state satisfies $\mathbf{Kq} = 0$.

The general three-state model allows direct transitions between any two states ($k_{ij} > 0$ for all $i$ and $j$). However, most gene regulation models to date followed the thermodynamic formalism with a detailed balance between states[27]. This constraint limits the topology of the state-transition diagram, i.e., transitions between certain states may be forbidden. Specifically, a three-state model with detailed balance needs to satisfy one of the two schemes of promoter activation, i.e., the sequential activation scheme, in which transitions between states 0 and 2 are forbidden, and the parallel activation scheme, where transitions between states 1 and 2 are not allowed (Supplementary Fig. 10a).

#### *6.4 Mean nascent mRNA signal*

The mean signal of the nascent mRNA may be derived from Equation (1) as follows[20]:

$$\langle m \rangle = \mathbf{u}\left\{\int_{-T_{\mathrm{RES}}}^{0} g(\tau)\mathbf{W}(\tau)d\tau\right\}\mathbf{q} \tag{2}$$

where $\mathbf{u} = (1, 1, 1)$ and $\mathbf{W}(\tau) = e^{-\mathbf{K}\tau}\mathbf{K}_{\mathrm{INI}}e^{\mathbf{K}\tau}$. At steady state, the magnitude of $\langle m \rangle$ is proportional to the mean of the contribution function, i.e.,

$$\langle m \rangle = \mathbf{u}\mathbf{K}_{\mathrm{INI}}\mathbf{q}\int_{-T_{\mathrm{RES}}}^{0} g(\tau)d\tau \tag{3}$$

#### *6.5 Estimating the post-elongation residence time*

According to Equation (3), the mean nascent mRNA signals measured using different probe sets are in proportion, i.e.

$$\frac{\langle m_1 \rangle}{\langle m_2 \rangle} = \frac{\overline{g_1}}{\overline{g_2}} \tag{4}$$

where $\overline{\cdot} = \frac{1}{T_{\mathrm{RES}}}\int_{-T_{\mathrm{RES}}}^{0} \cdot \, d\tau$ denotes time averaging. A probe set targeting the 5' region of a transcript should, on average, produce more signal (in units of the number of mRNA molecules) than a probe set targeting the 3' region of the same transcript. In our study, the ratios between the CDS and promoter-specific signals were defined as $a_1$ and $a_2$ in Equation (1). The target positions of these probe sets in mRNA sequences suggest $a_1 < 1$ and $a_2 > 1$, which agree with experimental results (Fig. 2c).

Notice that the time average of a contribution function can be divided into two parts, i.e.,

$$\bar{g} = \frac{1}{T_{\text{RES}}}\left(T_{\text{R}} + \int_{-L/V_{\text{EL}}}^{0} g(\tau)d\tau\right) = \frac{1}{T_{\text{RES}}}\left(T_{\text{R}} + \frac{L}{V_{\text{EL}}}\overline{g_0}\right) \tag{5}$$

The second term in parentheses may be defined as the time integration of a null contribution function ($g_0$) of the same probe set with $T_R = 0$. Since $g_0$ is independent of $T_R$ and only related to the target positions of the probe set, Equation (5) can be rewritten as

$$\frac{\langle m_1 \rangle}{\langle m_2 \rangle} = \frac{\overline{g_{10}} \cdot L + T_{\text{R}} V_{\text{EL}}}{\overline{g_{20}} \cdot L + T_{\text{R}} V_{\text{EL}}} \tag{6}$$

Or

$$T_{\text{R}} = \frac{L(\overline{g_{10}} - a\overline{g_{20}})}{V_{\text{EL}}\left(a-1\right)} \tag{7}$$

where $a = \frac{\langle m_1 \rangle}{\langle m_2 \rangle}$ is experimentally measurable.

Equation (7) may be used to estimate $T_R$ for P1- and P2-driven transcripts. For P1-driven transcripts, the null contribution functions of the CDS and P1-5'UTR probes satisfy $\overline{g_{\text{P1-CDS,0}}} = 0.2613$ and $\overline{g_{\text{P1-5'UTR,0}}} = 0.9613$, while the ratio between their nascent signals was measured to be $a_1 = 0.53$. Thus, with $L = 6334$ bp and $V_{EL} = 1.5$ kb/min[28], we estimated $T_R = 142$ s. For P2-driven transcripts, the null contribution functions of the CDS and P2-3'UTR probes satisfy $\overline{g_{\text{P2-CDS,0}}} = 0.5977$ and $\overline{g_{\text{P2-3'UTR,0}}} = 0.0514$, while the ratio between their nascent signals was measured to be $a_2 = 3.72$. With $L = 3621$ bp and $V_{EL} = 1.5$ kb/min[28], we estimated $T_R = 22$ s.

#### *6.6 Estimating the splicing rate*

Unlike probes targeting the exon or UTR regions of a transcript, the intron probe signal is affected by co-transcriptional splicing (Fig. 2d). Assuming a Poissonian slicing process occurring after the completion of intron synthesis with specific rate $k_{\text{splicing}}$, we wrote the average nascent intron signal per locus as

$$\langle m \rangle = \mathbf{u}\mathbf{K}_{\text{INI}}\mathbf{q}\int_{-T_{\text{RES}}}^{0} g(\tau)s(\tau)d\tau \tag{8}$$

where $s(\tau)$ is the intron survival probability (without being spliced) for a nascent transcript initiated at time $\tau$. $s(\tau)$ is a simple piecewise function satisfying

$$s(\tau) = \begin{cases} 1, & \tau > -L_{\text{5'-intron}} / V_{\text{EL}} \\ e^{k_{\text{splicing}}(\tau + T_{\text{5'-intron}})}, & \tau \le -L_{\text{5'-intron}} / V_{\text{EL}} \end{cases} \tag{9}$$

with $L_{\text{5'-intron}}$ denoting the sequence length from the 5' cap to the end of the intron. Thus, the mean nascent P1-intron and 5'UTR signals should be in proportion, with the ratio depending on $k_{\text{splicing}}$. The experimental data confirmed this linear relationship and suggested a ratio is of 0.59 (Supplementary Fig. 5d). Assuming that $V_{\text{EL}}$ = 1.5 kb/min[28] and $T_{\text{R-P1}}$ = 142 s, we estimated that $k_{\text{splicing}}$ = 175 s$^{-1}$. The time scale is similar to that observed in other genes[29-32].

#### *6.7 Variance and noise*

The variance of the nascent mRNA signal was derived from Equation (1) as[20]:

$$\sigma_m^2 = \mathbf{u} \cdot \left\{ \int_{-T_{\text{RES}}}^{0} d\tau_1 g(\tau_1)^2 \mathbf{W}(\tau_1) \right\} \mathbf{q}$$
$$+2\mathbf{u} \cdot \left\{ \int_{-T_{\text{RES}}}^{0} d\tau_1 \int_{-T_{\text{RES}}}^{\tau_1} d\tau_2 g(\tau_1) g(\tau_2) \mathbf{K}_{\text{INI}} \left( e^{\mathbf{K}(\tau_1 - \tau_2)} - \mathbf{qu} \right) \mathbf{K}_{\text{INI}} \right\} \mathbf{q} \tag{10}$$

In case of slow gene-state transitions, $e^{\mathbf{K}(\tau_1-\tau_2)} \approx \mathbf{I}$. Thus,

$$\sigma_m^2 \approx \mathbf{uK}_{\text{INI}}\mathbf{q} \int_{-T_{\text{RES}}}^{0} g(\tau)^2 d\tau + \mathbf{uK}_{\text{INI}} \left( \mathbf{I} - \mathbf{qu} \right) \mathbf{K}_{\text{INI}} \mathbf{q} \left( \int_{-T_{\text{RES}}}^{0} g(\tau) d\tau \right)^2 \tag{11}$$

Combining Equations (3) and (11), we wrote the noise of the nascent mRNA signal as

$$\eta^2 = \frac{\sigma_m^2}{\langle m \rangle^2} = \frac{1}{\langle m \rangle} \frac{\overline{g^2}}{\overline{g}} + \frac{\mathbf{uK}_{\text{INI}} \left( \mathbf{I} - \mathbf{qu} \right) \mathbf{K}_{\text{INI}} \mathbf{q}}{\left( \mathbf{uK}_{\text{INI}} \mathbf{q} \right)^2} \tag{12}$$

The first term in Equation (12) indicates Poisson noise, which is inversely proportional to $\langle m \rangle$. Its magnitude varies with the shape of *g*. The second term in Equation (12) is due to bursty expression, and the magnitude is invariant with the shape of *g*. For brevity, we rewrote the expression of noise as

$$\eta^2 = \frac{S_g}{\langle m \rangle} + \eta_{\text{burst}}^2 \tag{13}$$

where $s_g = \overline{g^2} / \overline{g}$ is a constant for a given probe set and mRNA species. For P1-driven transcripts, we had $S_{\text{P1-CDS}}$ = 0.78 for the CDS probes and $S_{\text{P1-5'UTR}}$ = 0.99 for the 5'UTR probes.

#### *6.8 Numerically solving the master equation*

Because the analytical solution for Equation (1) is not available, we solved the equation numerically using the finite state projection (FSP) method[26,33,34]. Briefly, we discretized and truncated the range of nascent mRNA signal to $m = 0, \Delta m, 2\Delta m, \ldots, m_{\max}$, with $\Delta m \ll 1$ and $m_{\max}$ large enough to cover the main portion of the nascent mRNA distribution. Equation (1) then transforms to a finite-dimension version:

$$\dot{\bar{\mathbf{P}}} = \left(\bar{\mathbf{K}} + \bar{\mathbf{K}}_{\mathrm{INI}}(\tau)\right)\bar{\mathbf{P}} = \begin{bmatrix} \mathbf{K}-\mathbf{K}_{\mathrm{INI}} & 0 & 0 & 0 \\ 0 & \mathbf{K}-\mathbf{K}_{\mathrm{INI}} & 0 & \cdots \\ 0 & 0 & \mathbf{K}-\mathbf{K}_{\mathrm{INI}} & \cdots \\ \vdots & \vdots & \vdots & \ddots \\ \mathbf{K}_{\mathrm{INI}} & 0 & 0 & \ddots \\ 0 & \mathbf{K}_{\mathrm{INI}} & 0 & \ddots \\ 0 & 0 & \mathbf{K}_{\mathrm{INI}} & \ddots \\ \vdots & \vdots & \vdots & \ddots \end{bmatrix} \begin{bmatrix} \mathbf{P}(0) \\ \mathbf{P}(\Delta m) \\ \mathbf{P}(2\Delta m) \\ \vdots \\ \mathbf{P}(g(\tau)) \\ \mathbf{P}(g(\tau)+\Delta m) \\ \mathbf{P}(g(\tau)+2\Delta m) \\ \vdots \end{bmatrix}, \tag{14}$$

where $\bar{\mathbf{K}} = \begin{bmatrix} \mathbf{K} & 0 & 0 & \cdots \\ 0 & \mathbf{K} & 0 & \cdots \\ 0 & 0 & \mathbf{K} & \cdots \\ \vdots & \vdots & \vdots & \ddots \end{bmatrix}$, $\bar{\mathbf{K}}_{\mathrm{INI}}(\tau) = \begin{bmatrix} -\mathbf{K}_{\mathrm{INI}} & 0 & 0 & \cdots \\ 0 & -\mathbf{K}_{\mathrm{INI}} & 0 & \cdots \\ \vdots & 0 & -\mathbf{K}_{\mathrm{INI}} & \cdots \\ \mathbf{K}_{\mathrm{INI}} & \vdots & 0 & \cdots \\ 0 & \mathbf{K}_{\mathrm{INI}} & \vdots & \ddots \\ 0 & 0 & \mathbf{K}_{\mathrm{INI}} & \ddots \\ \vdots & \vdots & \vdots & \ddots \end{bmatrix}$.

Next, we discretized the time range $\tau \in [-T_{\mathrm{RES}}, 0]$ into a series with $\Delta\tau \ll T_{\mathrm{RES}}$. The probability distribution of $(n, m)$ at $\tau = 0$ was computed by propagating the initial state $\bar{\mathbf{P}}_{\tau=-T_{\mathrm{RES}}}$ through the series, i.e.,

$$\bar{\mathbf{P}}_{\tau=0} = \left(\mathbf{I} + \bar{\mathbf{K}}\Delta\tau + \bar{\mathbf{K}}_{\mathrm{INI}}(-\Delta\tau)\Delta\tau\right)\cdots\left(\mathbf{I} + \bar{\mathbf{K}}\Delta\tau + \bar{\mathbf{K}}_{\mathrm{INI}}(-T_{\mathrm{RES}})\Delta\tau\right)\bar{\mathbf{P}}_{\tau=-T_{\mathrm{RES}}}, \tag{15}$$

where **I** is the unit matrix. In this paper, we used $\Delta m = 0.1$ and $\Delta \tau = T_{RES}/2000$ to balance the accuracy and speed of computation.

#### *6.9 Modeling the DNA replication effect*

The fact that some anterior nuclei contain more than two bright FISH spots (Supplementary Fig. 3a) indicates that the *hb* gene in the imaged embryo may have been replicated. Thus, many of the observed bright FISH spots may indeed be a pair of closely located sister loci that are indistinguishable under the microscope[22,23]. To consider this effect in the model/analysis, we note that the two sister gene copies are expressed independently[22,23]. The distribution of the observed signal from a pair of indistinguishable sister loci $P(m_{ob})$ should be a convolution of that of individual ones, i.e.,

$$P(m_{ob}) = P(m_{single}) * P(m_{single}) \tag{16}$$

where $P(m_{single})$ denotes the nascent mRNA distribution of a single gene copy computed from the model.

In general, we assume that a certain percentage ($\alpha$) of the observed FISH spots are from indistinguishable sister loci pairs. The total distribution was thus written as

$$P(m_{ob}) = (1-\alpha)P(m_{single}) + \alpha P(m_{single}) * P(m_{single}) \tag{17}$$

where $\alpha$ was determined by fitting $P(m_{ob})$ to experimental data (see Supplementary Note 6.10).

In addition to the probability distribution, the low-order statistics of the observed bright FISH spot are also affected by gene replication. Specifically, the mean and variance double with gene replication, while the Fano factor and correlation coefficient stay unchanged.

***6.10 Inferring the transcription kinetics***

We fitted the experimental data to estimate the kinetic parameters of each promoter using the maximum likelihood estimation (MLE) method[26,34]. Briefly, we divided the single-locus data of nascent mRNAs from an embryo into multiple subsets according to the nuclear position. To ensure a sufficient number of data points in each subset, we used overlapping binning with a bin size of 0.1 EL. For a given parameter set $\tilde{\mathbf{K}} = \{k_{ij}, k_{\mathrm{INI},i}\}$ and $\alpha$ (see Supplementary Note 6.9), the likelihood of observing a subset of data is

$$L(M \mid \alpha, \tilde{\mathbf{K}}) = \prod_i P(m_i \mid \alpha, \tilde{\mathbf{K}}) \tag{18}$$

where $P(m_i \mid \alpha, \tilde{\mathbf{K}})$ is the probability of observing $m_i$ nascent mRNAs given $\alpha$ and $\tilde{\mathbf{K}}$. For each subset $m_i$, we searched $\alpha$ and $\tilde{\mathbf{K}}$ to maximize the likelihood in a broad range of parameter values ($\alpha$ from 0 to 1, $k_{ij}$ from 0 to 10 min$^{-1}$, $k_{\mathrm{INI},i}$ from 0 to 100 min$^{-1}$).

To increase the efficiency and robustness of the parameter search for a three-state model, we first fitted a data set pooled from multiple embryos in the same nuclear cycle. For each nuclear position bin, we compared two types of models with either sequential or parallel activation schemes. Using a combination of simplex and simulated annealing methods for the parameter search, we determined that both P1 and P2 data were better fitted by the sequential activation model for all nuclear position bins. Moreover, the results showed that Bcd mainly affected promoter activation rates, while the inactivation and transcription initiation rates remained stable (Supplementary Fig. 10b). Thus, we fixed promoter inactivation and transcription initiation rates at their mean values and re-scanned the activation rates in detail. Besides $k_{ij}$ and $k_{\mathrm{INI},i}$, $\alpha$ was estimated to be ~0.5, a reasonable value considering the variation in the developmental time of different embryos. Once all kinetic rates were determined for the pooled data set, we applied them as initial values to fit the single-embryo data (except for the nc11 embryo, whose number of data points (<50 active loci per position bin of a single embryo) is too small for robust fitting). To increase the accuracy of simplex and simulated annealing methods in the above steps, we repeated each search 12 times. The result with the highest likelihood was chosen.

#### *6.11 Describing P1 and P2 activities using a single model*

Our results showed that P1 and P2 followed common three-state transcription kinetics driven by the same set of Bcd binding events at the two enhancers. Thus, we can combine the description of the two promoters into a single model to relate Bcd binding configurations with P1 and P2 transcription kinetics (Fig. 5i).

The first part of the model describes the Bcd binding dynamics. There are many Bcd binding sites on the proximal and distal enhancers[35-37]. For simplicity, we assumed that Bcd binding at each enhancer was highly cooperative, with all binding sites being occupied/emptied in one step. This assumption resulted in four possible Bcd binding configurations (Fig. 5i). In the canonical framework of transcription factor binding dynamics, transitions between these binding configurations are described as Poisson processes, whose kinetic rates are related to Bcd concentration by a power law[38]. The steady-state probability of each binding configuration (*s*) satisfies a rational function,

$$P_s(C_{\mathrm{Bcd}}) = \frac{r_s C_{\mathrm{Bcd}}^{n_s}}{\sum_s r_s C_{\mathrm{Bcd}}^{n_s}} \tag{19}$$

where $C_{\mathrm{Bcd}}$ is the Bcd concentration, $n_s$ and $r_s$ are the power-law exponents and proportionality constants for configuration *s*, respectively. Specifically, the configuration with no Bcd bound at either enhancer, (typically denoted as $s = 0$) satisfies $n_0 = 0$ and $r_0 = 1$. For an equilibrium system satisfying detailed balance[27], $n_s$ equals the number of bound Bcd molecules. For a nonequilibrium system, $n_s$ may take higher values[38], yet the general form of Equation (19) still holds.

To relate Bcd binding with the transcriptional activity of a promoter, we assumed that transitions between different promoter states were triggered by specific Bcd binding configurations (Fig. 5i). Bcd binding at a single enhancer (proximal or distal) triggers the transition of a promoter from state 0 to state 1, while the binding at both enhancers triggers the transition from state 1 to state 2. Strictly speaking, these transitions can only happen when the system is at given Bcd binding configurations. However, since transcription factor binding and unbinding happen at a much faster time scale than promoter activation[23], the promoter activation rates ($k_{01}$

and $k_{12}$) can be modeled as constants over time.

In a simple model, the promoter activation rate may be proportional to the probability of the corresponding Bcd binding configuration. However, activation of a real promoter involves a series of molecular events, some of which are independent of Bcd[23,26]. The activation rate estimated from nascent mRNA distribution ($k_{01}$ or $k_{12}$) represents the overall time scale of all molecular events, i.e.,

$$\begin{cases} k_{01}^{-1} = \left[a_1 P_1(C_{\mathrm{Bcd}}) + a_2 P_2(C_{\mathrm{Bcd}})\right]^{-1} + \tau_{01} \\ k_{12}^{-1} = \left[b P_3(C_{\mathrm{Bcd}})\right]^{-1} + \tau_{12} \end{cases} \tag{20}$$

where $a$ and $b$ are proportionality constants and $s$ = 1, 2, 3 denote the binding configurations with Bcd bound at the proximal, distal, or both enhancers, respectively. $\tau$ represents the time scale of Bcd-independent molecular events, which can saturate $k_{01}$ and $k_{12}$ at high Bcd concentration. Equation (20) explains the Hill-function-like relationship between promoter activation rates ($k_{01}$ and $k_{12}$) and Bcd concentration observed in Fig. 5c.

Since P1 and P2 nascent mRNA signals have little correlation (Supplementary Fig. 3d), we speculated that the activation of the two promoters was triggered independently. Thus, the joint distribution of P1 and P2 nascent mRNA signals is the product of their marginal distributions (Fig. 5j).

### Supplementary Figures

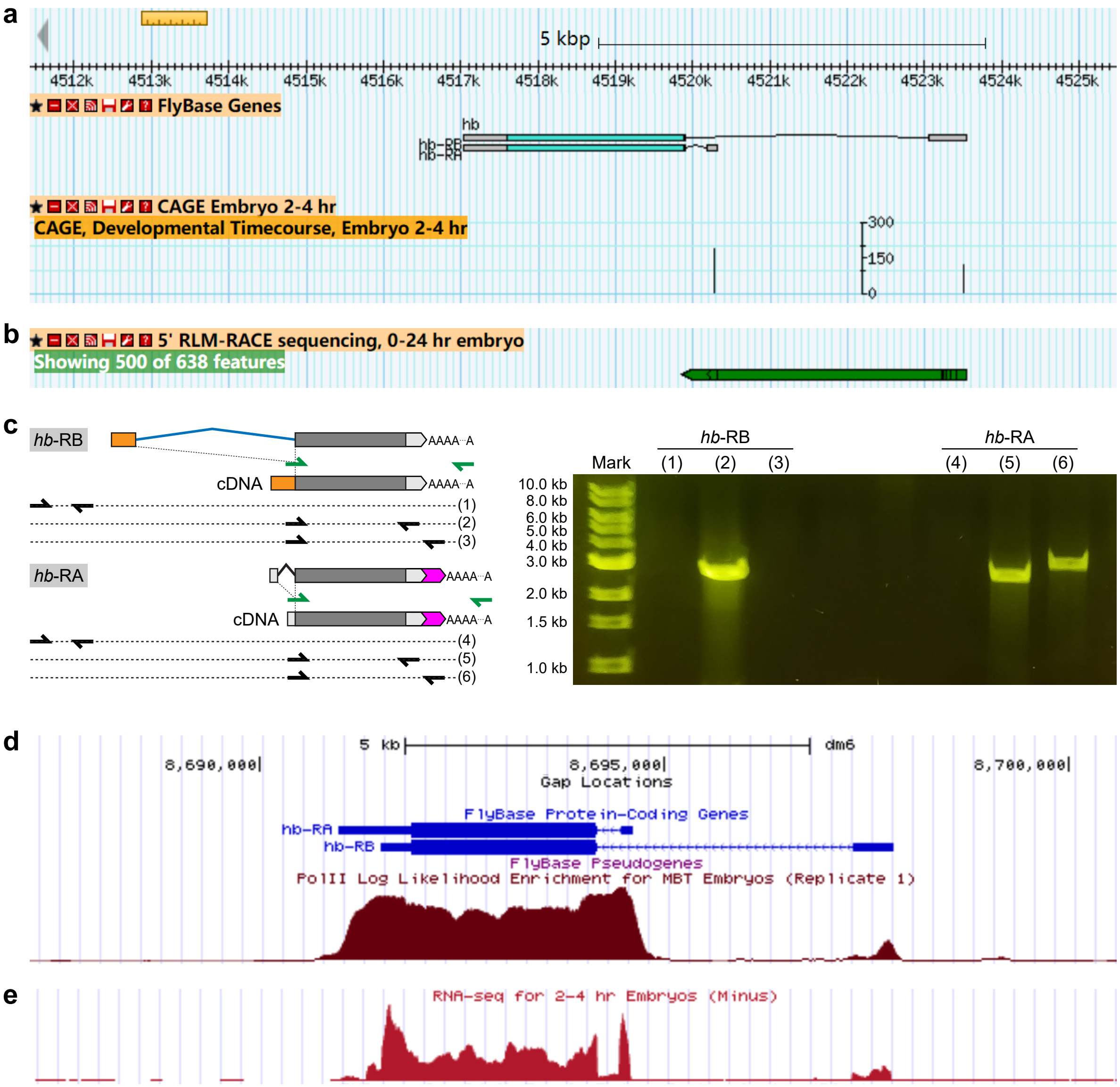


**Supplementary Figure 1. Bulk measurements of early *hb* promoter activity. (a, b)** Published CAGE **(a)** and 5'RACE **(b)** signals of *hb* in the developing embryo. Data from the Berkeley Drosophila Transcription Network Project were plotted using the Generic Genome Browser (http://gbrowse.modencode.org/). Both signals showed that *hb* mRNAs in early embryos are transcribed from two starting sites corresponding to P1 and P2, respectively. **(c)** 3'RACE and PCR experiments to verify the 3' ends of P1- and P2-driven transcripts. Left: schematic of 3'RACE and PCR (see **Supplementary Note 1**). Right: agarose gel electrophoresis of PCR products. P2-driven transcripts terminate at a more distant site than P1-driven transcripts. **(e, f)** Published PolII ChIP-seq **(e)** and RNA-seq **(f)** signals of *hb* in the developing embryo. Data were from the Gene Expression Omnibus database at NCBI (accession number: GSE41700) and the ENCODE data portal at UCSC, respectively. Data were plotted using the GEP UCSC Genome Browser (https://gander.wustl.edu/). Both signals exhibit peaks at P1 and P2 promoters. The peak at P1 is much lower than that at P2, suggesting that P1 is less active than P2 in the early embryo.

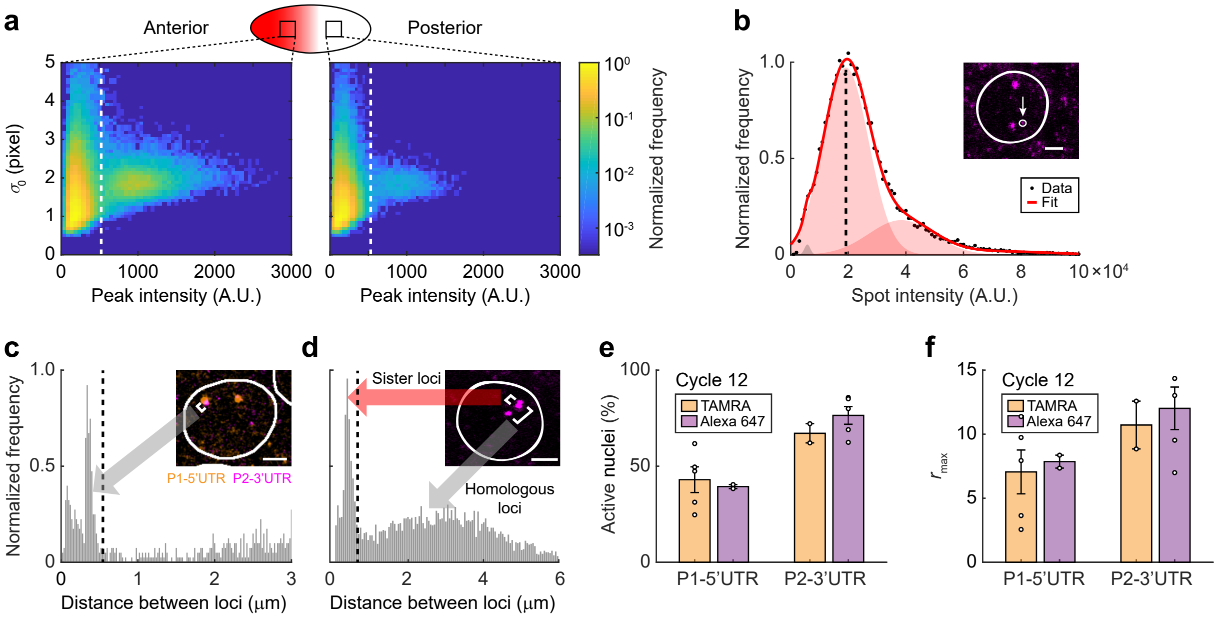


**Supplementary Figure 2. Quantifying *hb* mRNA. (a)** The joint distribution of peak height and radius for candidate smFISH spots in the anterior (left) and posterior (right) parts of the embryo (>60,000 CDS spots from a single nc12 embryo). A threshold (dashed line) was used to identify spots corresponding to real *hb* mRNAs from false-positive particles. **(b)** The intensity histogram of smFISH spots at the anterior part of the embryo (>20,000 spots). The intensity corresponding to a single mRNA molecule was identified by fitting the histogram to the sum of Gaussian functions. Inset: smFISH signal in the anterior part of the embryo. Scale bar, 2 μm. **(c)** The distribution of the distance between smFISH spots from P1-5'UTR and P2-3'UTR channels (>30,000 spots in a single nc12 embryo). Two groups of spot pairs were recognized from the distribution. A distance threshold was used to identify inter-channel spot pairs corresponding to the same transcription site. Inset: P1-5'UTR and P2-3'UTR signals in an anterior nucleus. Scale bar, 2 μm. **(d)** The distribution of the distance between smFISH spots in the same nucleus (>30,000 P2-3'UTR spots in a single nc12 embryo). Two groups of spot pairs were recognized from the distribution. A distance threshold was used to identify sister loci pairs. Inset: smFISH signal in an anterior nucleus. Scale bar, 2 μm. **(e, f)** The average percentage of active nuclei **(e)** and the maximal nuclear signal level **(f)** of P1 and P2 expression domains were measured using two groups of P1-5'UTR and P2-3'UTR probe with switched fluorophores (TAMRA and Alexa Fluor™ 647). Neither quantity changed significantly with fluorophore switching, indicating that the efficiencies of different fluorescent detectors are comparable. Data averaged from ≥2 nc12 embryos for each group of probe sets. Error bars represent s.e.m.

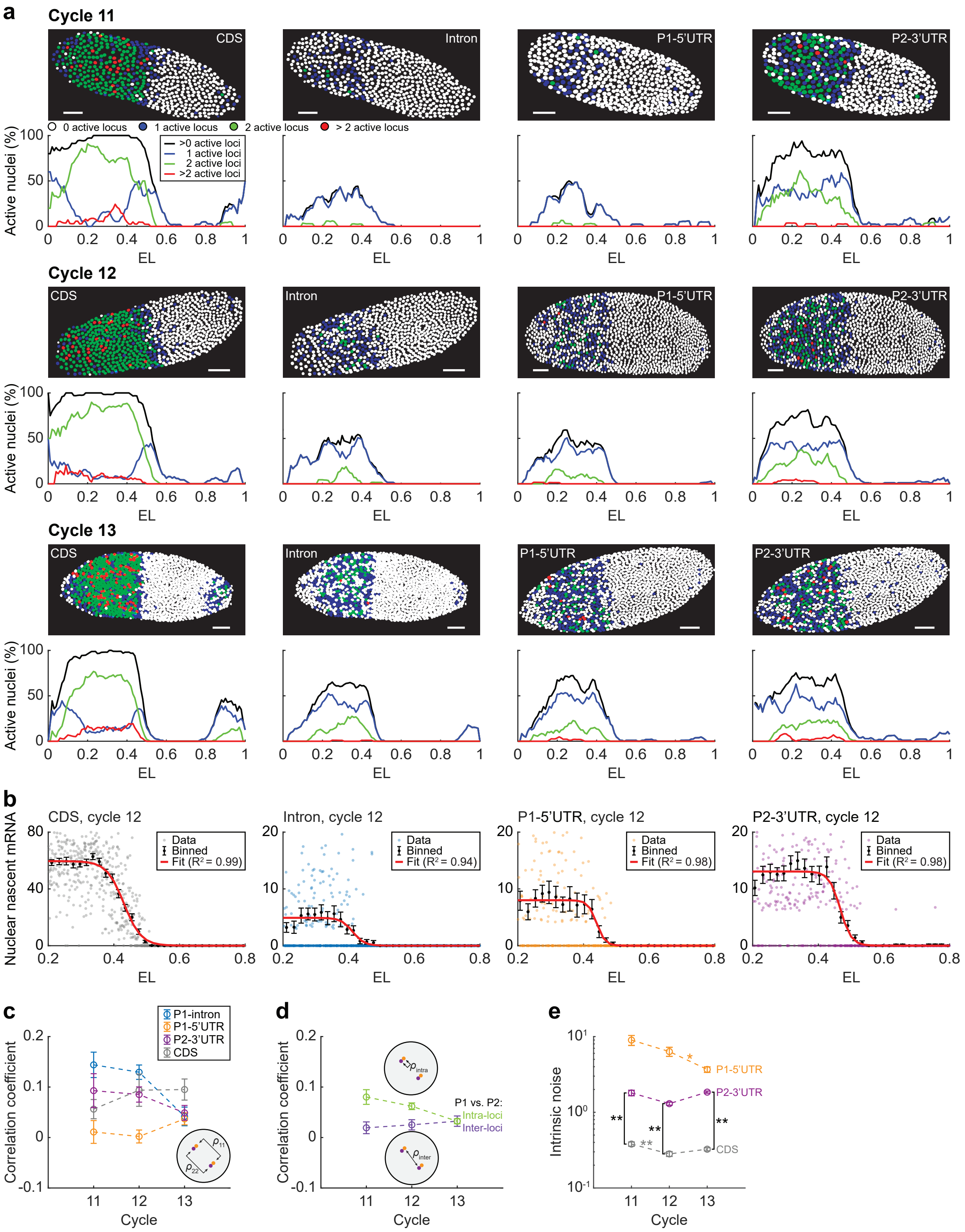


**Supplementary Figure 3. The expression profile and fluctuation of the two *hb* promoters. (a)** The transcriptional activity of individual nuclei in WT embryos during nc11–13 measured using different probe sets. The color of each nucleus indicates the number of active *hb* loci (see legend). Scale bar, 50 μm. The percentage of nuclei containing different numbers of active loci as a function of the AP position was attached below. **(b)** Nascent signal in individual nuclei (in units of the number of mRNA molecules) was plotted against the AP position for different probe sets from individual embryos. The single-nucleus data were binned along the AP axis (mean ± s.e.m.) and fitted to Logistic functions. **(c)** The correlation coefficient of the nascent mRNA signal between different *hb* gene loci in the same nucleus in the position range of 0.2–0.4 EL for different probe sets during nc11–13. **(d)** The correlation coefficient between P1-5'UTR and P2-3'UTR signals from the same (intra-allele) or different (inter-allele) *hb* gene loci in the same nucleus in the position range of 0.2–0.4 EL during nc11–13. **(e)** The intrinsic noise of P1-5'UTR, P2-3'UTR, and CDS signals at individual *hb* gene loci in the position range of 0.2–0.4 EL during nc11–13. The noise of P1-5'UTR was 2–5 times higher than that of P2-3'UTR. This was mainly due to the low expression level of P1. Since the CDS signal is the sum of P1 and P2 activities, its noise level is significantly lower than P2-3'UTR by ~5 folds. P-values were from Student's t-test: *, $p<0.05$; **, $p<0.01$. **(a–b)** The single-nucleus data were binned along the AP axis (bin size: 0.05 EL, overlap between neighboring bins: 50%). **(c–e)** Data averaged from ≥ 5 embryos for each nuclear cycle. Error bars represent s.e.m.

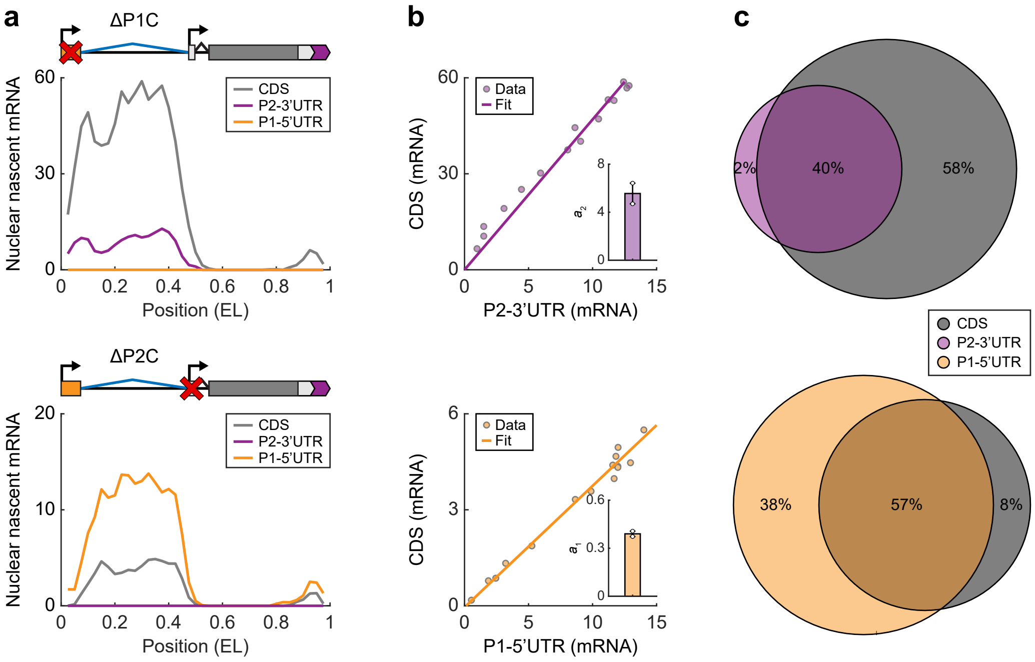


**Supplementary Figure 4. The relationship between different probe signals in promoter deletion embryos. (a)** Average nascent P1-5'UTR, P2-3'UTR, and CDS signals per nucleus (in units of the number of molecules) as functions of the AP position in individual embryos with P1 or P2 deletion (bin size: 0.05 EL, step size: 0.025 EL). **(b)** Nascent CDS and promoter-specific signals per nucleus at different AP positions (0.3–0.6 EL) were plotted against each other and fitted to a linear function for individual embryos with P1 or P2 deletion. Insets, the ratios between CDS and promoter specific probes. The values were roughly similar to $a_2$ and $a_1$ estimated from the WT embryo (See **Supplementary Note 3**). Data averaged from two embryos for each fly line. **(c)** Number and colocalization statistics of CDS and promoter-specific foci in promoter deletion embryos. The P1 deletion embryo contained more CDS foci than P2-3'UTR foci, while the P2 deletion embryo contained more P1-5'UTR foci than CDS foci. In both constructs, foci of 3' probes were mostly colocalized with that of 5' probes.

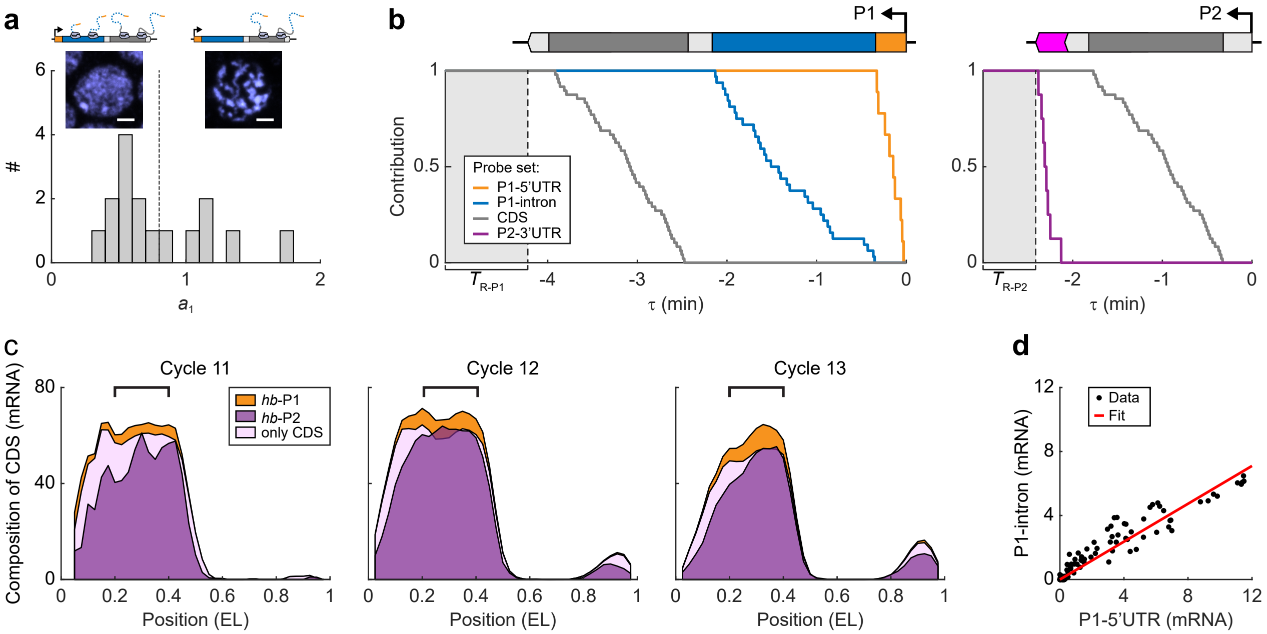


**Supplementary Figure 5. Decomposing nascent *hb* transcription into P1 and P2 activities. (a)** The distribution of $a_1$ extracted from 18 embryos in nc11–13. Embryos in mitotic interphase (nuclear morphology shown in the left inset, scale bar, 2 μm) exhibited $a_1 \sim 0.5$, while those close to mitosis (nuclear morphology shown in the right inset, scale bar, 2 μm) exhibited $a_1 \gtrsim 1$. The difference between them was due to the shut-down of transcription initiation at the end of the cell cycle (top schematics). A threshold $a_1$ (dashed line) was used to separate interphase and premitotic embryos. One nc11, five nc12, and four nc13 interphase embryos were selected for analysis. **(b)** The contribution functions of different probe sets for P1- and P2-driven transcripts. For each probe set, the observed smFISH signals of a nascent transcript were plotted against the transcript's initiation time. **(c)** Reconstructing the CDS expression profile from P1- and P2-specific signals for different nuclear cycles using a1 and a2. Data from embryos selected by **(a)**. Bin size: 0.05 EL, step size: 0.025 EL. Promoter-specific signals accounted for most of the CDS signal in the anterior expression domain (marked region, 0.2–0.4 EL). A fraction of the CDS signal in the terminal regions (0–0.2 and 0.8–1 EL) was not covered, suggesting that different regions of the embryo may have different $a_1$ (and $a_2$). **(d)** The average P1-5'UTR and P1-intron signals per nucleus at different AP positions (bin size: 0.05 EL, step size: 0.025 EL) were plotted against each other and fitted to a linear function. Data pooled from 15 embryos during nc11–13 for each probe signal.

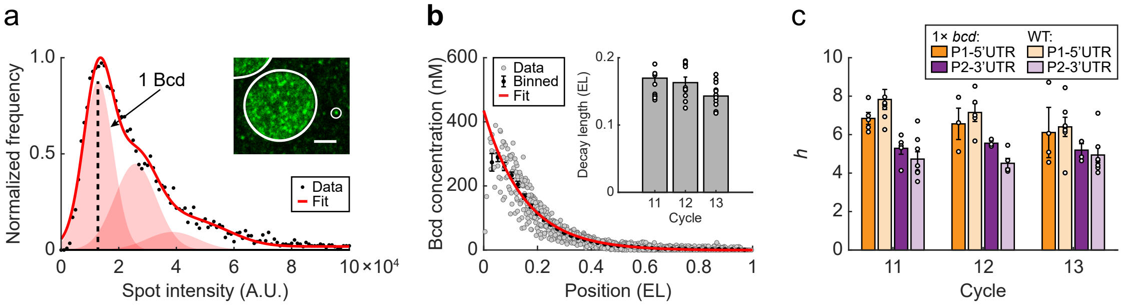


**Supplementary Figure 6. Quantifying Bcd concentration and its regulation of *hb* promoters. (a)** The intensity histogram of cytoplasmic Bcd spots at the anterior part of the embryo (>20,000 spots). The intensity corresponding to a single Bcd protein was identified by fitting the histogram to the sum of Gaussian functions. Inset: immunofluorescence signal of Bcd in the anterior part of the embryo. Scale bar, 2 μm. **(b)** The exponential gradient of nuclear Bcd concentration along the AP axis of a WT embryo in nc12. Data from individual nuclei were binned along the AP axis (mean ± s.e.m., bin size: 0.05 EL, step size: 0.025 EL) and fitted to an exponential function. Inset: the decay length of the Bcd gradient. Data averaged from ≥10 embryos for each nuclear cycle. Error bars represent s.e.m. **(c)** The Hill coefficient of the gene regulation function for P1-5'UTR and P2-3'UTR signals in WT and 1x *bcd* embryos during nc11–13. Data averaged from ≥5 WT embryos and ≥3 1x *bcd* embryos for each nuclear cycle. Error bars represent s.e.m.

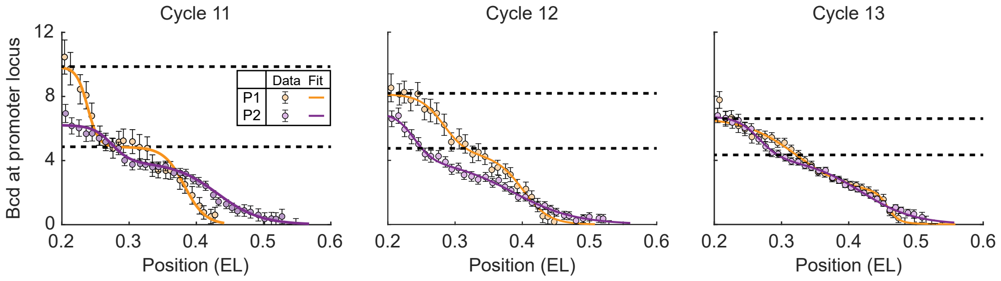


**Supplementary Figure 7. Quantifying Bcd binding at *hb* locus.** The average number of Bcd molecules bound at P1- and P2-active *hb* loci as a function of nuclear position during nc11–13. Data of each nuclear cycle were pooled from ≥5 embryos and binned along the AP axis (bin size: 0.05 EL, step size: 0.025 EL). The binned data were fitted to multi-logistic functions. Dashed lines highlight discrete binding plateaus for each promoter. In the very anterior part of the embryo (<0.25 EL), the P2-specific binding curve exhibited an additional plateau with ~6 Bcd molecules. In nc11–13, this plateau is lower than the P1-specific plateau that appeared in the same position range. Error bars represent s.e.m.

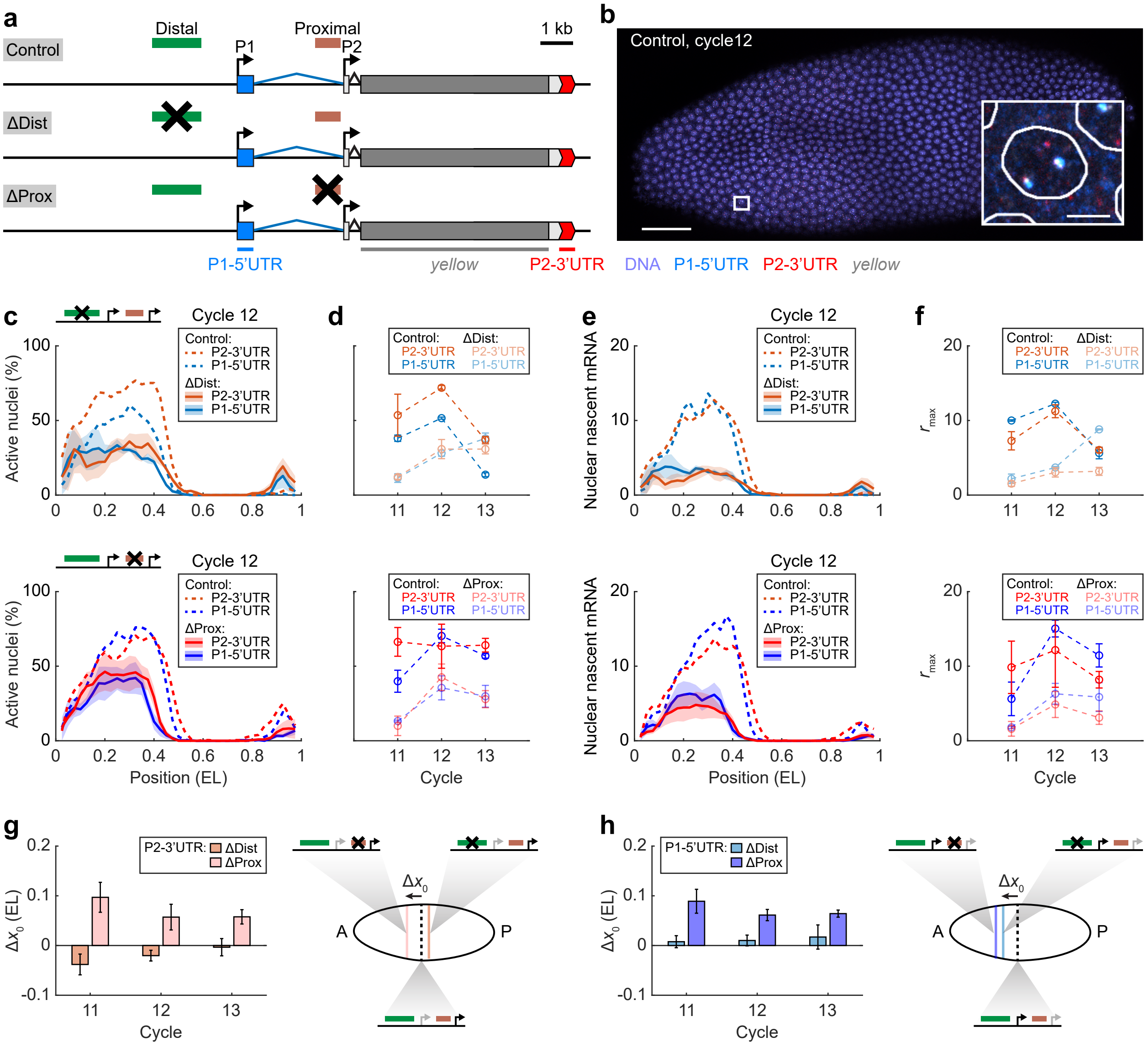


**Supplementary Figure 8. Measuring promoter activities in enhancer deletion constructs using P1-5'UTR and P2-3'UTR probes. (a)** Schematic of *hb* reporter constructs with enhancer replacements. Three smFISH probe sets were used to label different regions of reporter mRNAs: blue, P1-5'UTR probes; red, P2-3'UTR probes; grey, *yellow* probes. **(b)** Confocal image of a distal-enhancer-removed embryo labeled for P1-5'UTR, P2-3'UTR, and *yellow* at nc12. P1-5'UTR and P2-3'UTR signals corresponding to the transgene were identified based on their colocalization with *yellow* foci. Scale bar, 50 μm. Inset, magnified view of a single anterior nucleus. Scale bar, 2 μm. **(c)** Percentage of active nuclei as a function of the AP position for P1-5'UTR and P2-3'UTR signals in nc12 embryos of different constructs. Shadings indicate s.e.m. **(d)** The average percentage of active nuclei in the position range of 0.2–0.4 EL for P1-5'UTR and P2-3'UTR signals in different constructs during nc11–13. Error bars represent s.e.m. **(e)** Nascent P1-5'UTR and P2-3'UTR signals per nucleus as functions of the AP position in nc12 embryos of different constructs. Shadings indicate s.e.m. **(f)** The maximal signal level of the anterior expression domain for P1-5'UTR and P2-3'UTR signals in different constructs during nc11–13. Error bars were computed from the standard errors of boundary positions for enhancer-deleted and control lines. **(g, h)** The boundary shift of the anterior expression domain for P2-3'UTR **(g)** and P1-5'UTR **(h)** signals upon removing one enhancer. Error bars represent s.e.m. Right: schematic of boundary shift. **(c–h)** Data compared between the distal- or proximal-enhancer-removed constructs and their controls. Data averaged from ≥2 embryos for each reporter construct and each nuclear cycle. The spatial profile of each embryo was binned from the single-nucleus data (bin size: 0.05 EL, step size: 0.025 EL). P-values were from Student's t-test: *, $p<0.05$; **, $p<0.01$.

* In general, the results shown here are similar to **Fig. 4**. A slight difference between **Fig. 4g** and **Supplementary Fig. 8g** is on the magnitude of the boundary shift. This may be because the reporter gene is too long (~10 kbp) and does not have enough time to reach steady-state transcription in a nuclear cycle. Specifically, when the P1-5'UTR signal of the reporter gene reaches a high value, the P2-3'UTR signal may just emerge. When the P2-3'UTR signal reaches a high value, the P1-5'UTR signal may have diminished due to transcription shut down at the end of the nuclear cycle. In such a case, labeling the intron and CDS (*yellow*) regions may be a better choice, as they are relatively close to each other.

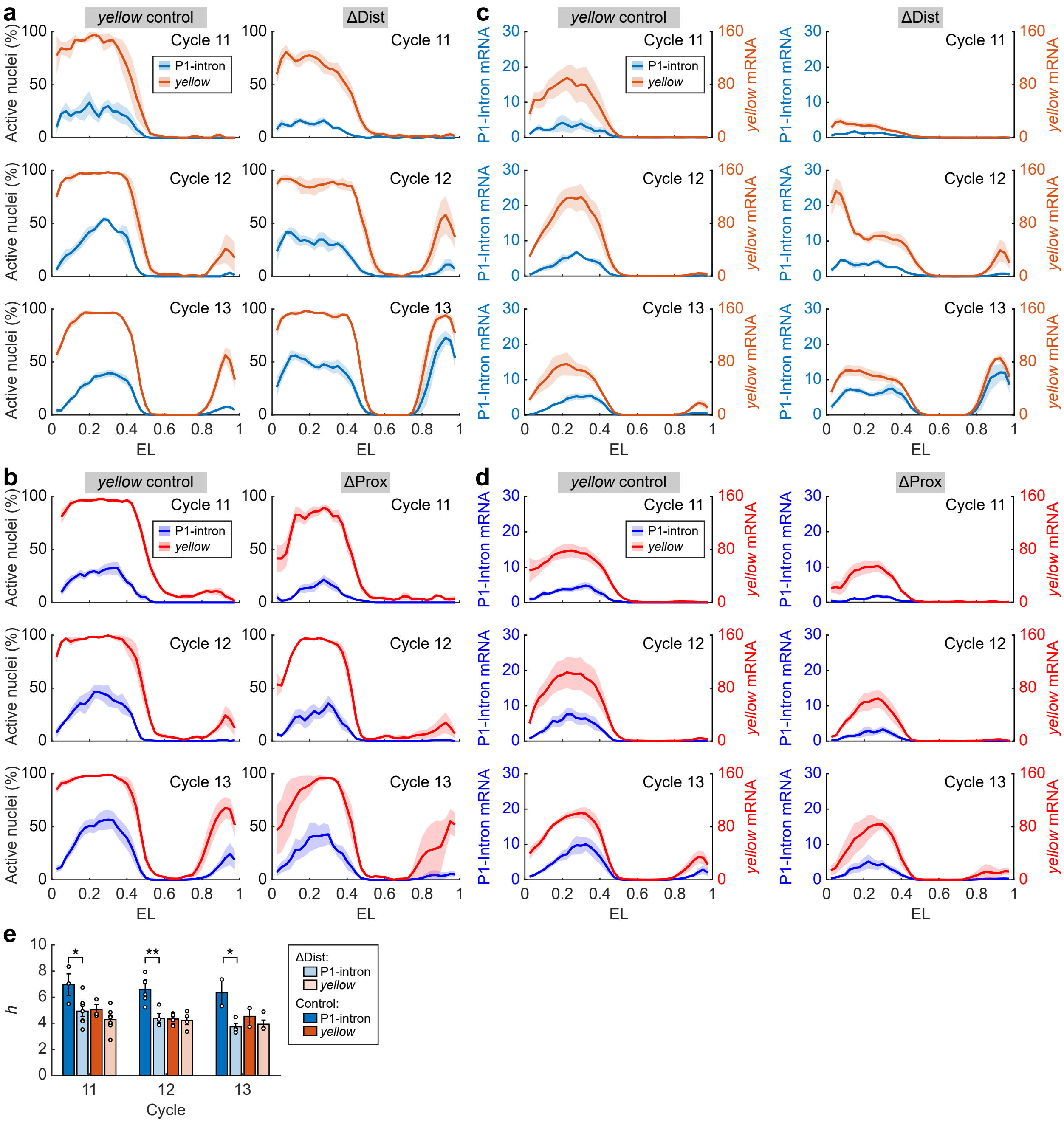


**Supplementary Figure 9. Comparing promoter activities in different reporter constructs. (a)** Percentage of active nuclei as a function of the AP position for P1-intron or *yellow* signals in the distal-enhancer-removed and control embryos. **(b)** Percentage of active nuclei as a function of the AP position for P1-intron or *yellow* signals in the proximal-enhancer-removed and control embryos. **(c)** Nascent P1-intron and *yellow* signals per nucleus as functions of the AP position in the distal-enhancer-removed and control embryos. **(d)** Nascent P1-intron and *yellow* signals per nucleus as functions of the AP position in the proximal-enhancer-removed and control embryos. **(e)** The Hill coefficient of the gene regulation function for P1-intron and *yellow* signals in the distal-enhancer-removed and control embryos during nc11–13. Data averaged from ≥2 embryos for each construct and each nuclear cycle. Error bars represent s.e.m. **(a, c)** Besides affecting the anterior expression domain, removing the distal enhancer also increased the expression of *yellow* and P1 in the terminal regions (0–0.2 and 0.8–1 EL). This observation agrees with a previous report that the distal enhancer may inhibit P1 and P2 at poles[37]. **(a–d)** Data averaged from ≥4 embryos for each construct and each nuclear cycle. The spatial profile of each embryo was binned from the single-nucleus data (bin size: 0.05 EL, step size: 0.025 EL). Shadings indicate s.e.m.

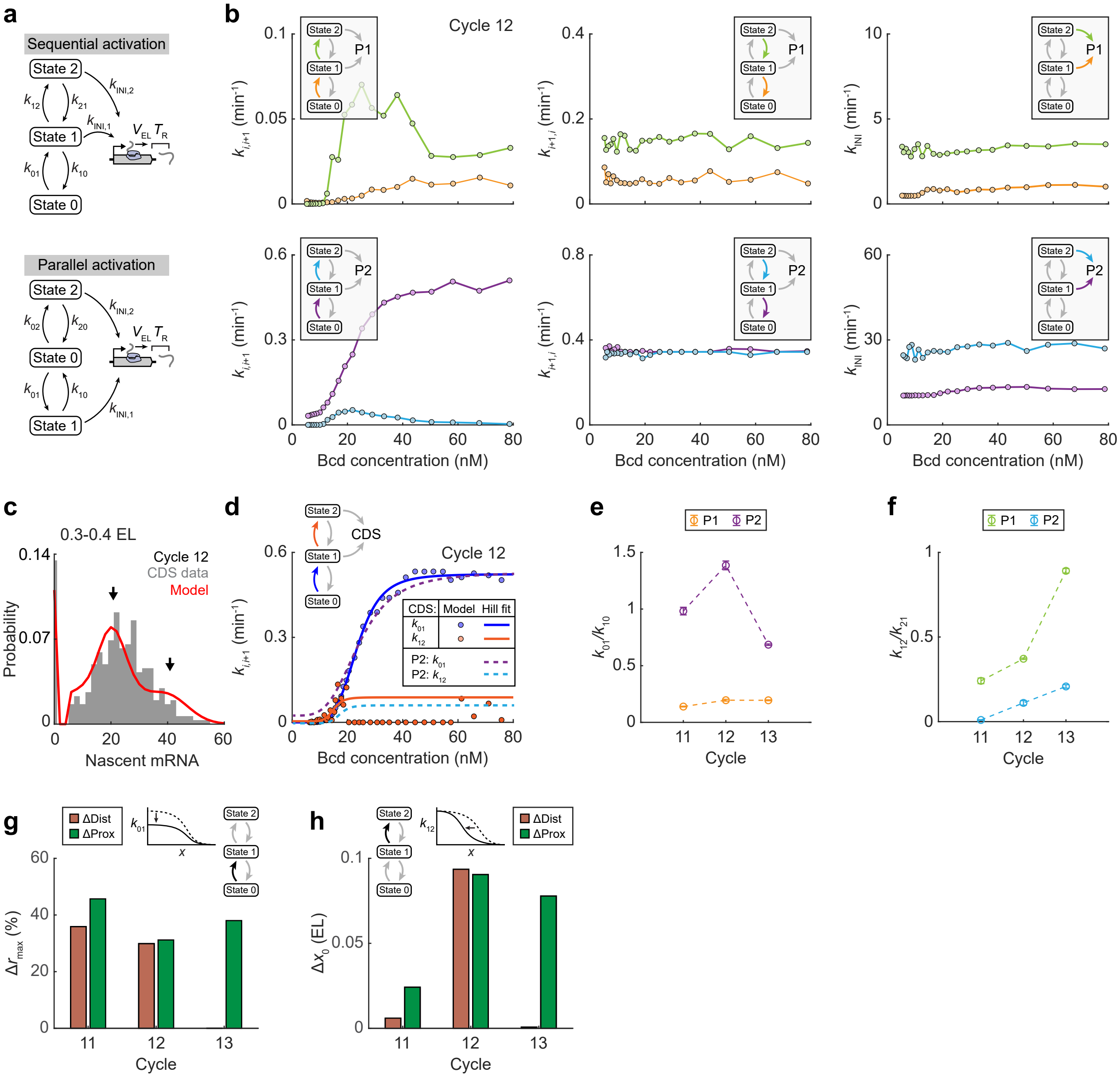

**Supplementary Figure 10. Fitting P1 and P2 nascent mRNA signals using a three-state model. (a)** Schematic of sequential and parallel activation schemes of the three-state model. In the sequential activation scheme, the promoter can be activated from state 0 to state 1 and from state 1 to state 2. In the parallel activation scheme, the promoter can be activated from state 0 to states 1 or 2. **(b)** The extracted parameters of the three-state model (binned by the AP position, bin size: 0.1 EL, step size: 0.01 EL) as a function of nuclear Bcd concentration. Data from five embryos at nc12. **(c)** Histogram of nascent CDS signals at individual *hb* gene loci in position range 0.3–0.4 EL of a single embryo. The histogram was fitted to a three-state transcription model. **(d)** The equivalent gene activation rates for the CDS signal estimated from two embryos at nc12 were plotted against nuclear Bcd concentration and fitted to Hill functions. The extracted $k_{01}$ and $k_{12}$ were similar to that from the P2-3'UTR signal, suggesting that CDS may be used as a proxy for P2. $k_{12}$ estimated here was a little bit bigger than that from the P2-3'UTR signal, revealing an influence from P1. **(e)** The average ratio between $k_{01}/k_{10}$ for P1 and P2 in the position range of 0.2–0.4 EL during nc11–13. Error bars represent s.e.m. **(f)** The average ratio between $k_{12}/k_{21}$ for P1 and P2 in the position range of 0.2–0.4 EL during nc11–13. Error bars represent s.e.m. **(g)** The relative decrease of the maximal $k_{01}$ level for P1 in the anterior expression domain upon removing one enhancer. In nc11–12, deleting either enhancer caused a decrease of the maximal $k_{01}$ level by ~30%–40%. In contrast, only the removal of the proximal enhancer caused a dramatic decrease of the maximal $k_{01}$ level in nc13. **(h)** The boundary shift of the anterior $k_{12}$ profile towards the anterior pole for P1 upon removing one enhancer. In nc11–12, deleting either enhancer caused an anterior shift of the $k_{12}$ expression boundary. The estimated boundary shift in nc11 is smaller than in nc12, probably because P1 is much less active in nc11 than in nc12–13 (**Fig. 2e**). In contrast, only the removal of the proximal enhancer caused a dramatic a dramatic anterior shift of the $k_{12}$ expression boundary in nc13. This is consistent with the decrease of the higher Bcd binding plateau in nc13 (**Supplementary Fig. 7**) and indicates a change in *hb* regulation. **(e, f)** Data averaged from ≥5 embryos for each nuclear cycle. **(g, h)** Data averaged from ≥4 embryos for each reporter construct and each nuclear cycle.

**Supplementary Table 1. Sequences of smFISH probes.**

| Target | Probe sequences (5’ to 3’) |
|---|---|
| P1-5’UTR | AATGCTGGCGACTTTCGTTT<br>TTTGTATTTCAGTGGCTGCC<br>CGTCCTTTGGATGTTTGGTT<br>CGGGACAAAAGTCTTTTTGC<br>GGATGTGGTCTTTGCCAAAA<br>TTTTGGGCCTCGCTTTTTAG<br>TTAGACCAACACGCACAGTG<br>GGGAGAATTTGGAAACGGGA<br>GGACAGTCCAAGTGCAATTC |
| P1-Intron | TAGGATATTGGATGGTACGC<br>GGAGGTCGAATGCAGTGTAT<br>CTCCCGCAAAAGCGATTTGT<br>GGCGTATGTAAATCCTCACA<br>TTGGATATACGCTGCAGTTC<br>CGTTGCTTAGGAGGAGCAAA<br>AATTAGCTGTCAGGCGTAGA<br>ATTGAAGGGGTTATTTGGGG<br>GTCGAAAATGCATTTGCCGA<br>AGCGTCACATAGGTGTATTC<br>CGGATGACAATCAATTCTGC<br>CCCATTTGTTTGAAATGCGC<br>AAAGCAAGCCAAGAGCAGGA<br>GCAGAAAATGCGCCACACAA<br>CATGTGCTAGTCGTTTTTGG<br>CAGGCAACCGAAACTGCAAA<br>GCCAAACTAAAGGCCAAGTG<br>TGTGTGTGCGCACTATGAAA<br>GCCAAAATTAATTGCTCGGC<br>GCACCACACAAAATGAAGCT<br>GGCATAATTGATGGTTCAGG<br>TTATTATGGGAGGATGGTGC<br>CGGGAAAAAGGGGCATTTAC<br>TGACAACAATTTTCCGCCAG<br>ACGGATCAGAACTGCTTACA<br>AGGATTGCGGGACTTAACTA<br>GGTTTTCTATGGGGATTACG<br>TAGCAGCGAGCTGCGAATTT<br>CGCACTTGGATTTGGATGAT<br>GATCCATTCTGGATTAGAGC<br>CACGCGTCAAGGGATTAGAT<br>TATATCGCTCAGGTAGACGG |
| CDS | TTGTGCTGCTCGTAGTTGGT<br>GAACATGCTGTTGTACCAGG<br>GCTCCTGTTTGATATTTGCC |

| | |
|---|---|
| | TATTCCCGTCGAGATGATGA<br>AACTGTTCCAGGTGATTGGT<br>ATCCATGGGTTGCTGCTGAA<br>TTTGATCGTTTTGGCTGGGT<br>TTAGCATCGTAATGCTGCAG<br>TTGCTGCAGCAACTGTTGCT<br>TGGAAATGCTGCTGGTACTG<br>ATGGTGATGATGTTGCTGCT<br>TTGAATCCACCCATCAGATG<br>TAGAAGTGCTGCATGGGATT<br>TGTTAGTGCCTGCAACTTCT<br>TATTCGACTGACTCGACTTG<br>ATGTACTTCATGTCCTCGCT<br>ATGTTGGTATCATCGTCCTC<br>TGCGAATTGTAGATGGGCAT<br>GGTCTTGCACTTGTAGTTCT<br>TTGTCTGGTTTCATGTGGGT<br>TACTCCAAGTGGTGCTTGAA<br>GTTCTTGTGCTTCCGGATAT<br>ACACGTGTAGCTGCATTTGT<br>GCGAGTTTAGCATGGATTTG<br>TACACAGAACTGTGCGACTT<br>TAATCACAATCCGCACAACG<br>AAGCTGTGGCAATACTTGGT<br>ATACTTGCGCAGATGCAGCT<br>AAACATCGATGACCAACGAG<br>ATTCTTGCTCTTCGGACCAC<br>AGCTGCAACATTTGACTTCC<br>TGGCTGAGATTGCTGTTGCT<br>TTGAACCAGAGGGAATCCTT<br>AAGAAGGCCATGTTGCGGTT<br>TGGAGATTGAGGTTCCAGTA<br>TCGCATTCTTGGCGACAATT<br>TGGTTCTGTTGCTGCAGTTG<br>TGACTTACGCTCGTACTCAT<br>TTCCTTGGGACAGATCCATG<br>TTGTTGCTGCTGCTCATCCT<br>TCCTCCACCTTGAGATTCAT<br>TGTGCTGGGTACTTTCAGTT<br>TTGCTATTGCTGCTGGCATT<br>TTCCATTGCTGCTGGAATTG<br>AGTACTTGCACTCGTAGATG<br>GCGTCCTTGAAGAAGATATC<br>CATGTGAATGGTGTAGAGCA<br>TTGCACTTGAACACATCGTC |
| P2-specific 3’UTR | AGAACTGAGTGTTATGCGCA<br>TCTTCTTTCGTCAGTTTCAG<br>CGCATCTTAGCTACTCTTTA |

| | |
|---|---|
| | AATTTTGATCCGTTGCTCAG<br>CGACTTAGATTTTATGGGGT<br>GTCTCGAAATTCGTTTCATG<br>ATCAAGGATTACACTGGGCT<br>CATTTCGTGGGCAAATATCT |
| *yellow* | AAACTGCGGTCCATGTTTAT<br>GCCAATCTGGATACGGAATT<br>CAATCTCCAGCTGTATTTGA<br>GTAGGCAGTGGTAATACTGT<br>CCACACTCATCCACTTTAAT<br>ACGGTTCCAGTGTCCAAAAC<br>CACGGATTAGTGGTGGTATT<br>GTATCCGTGGTCAAGTCAAA<br>TAGCTCGTATCTCCGAATTC<br>GTATTTGGATTTGTGTCCAC<br>CACGGCAATGTTAGCTATGA<br>TATCCCAATTCATCGGCAAA<br>CCCAGGAGTAAGCAATCAAG<br>AGAATCTCCAGGACTTGTTC<br>CCTCAATGGATCGGGGAAAA<br>CCCATTGGAAGTTAATACCA<br>ATACCAAATATACCCTCCTC<br>AGTACAGGGTACGATAACCA<br>CGATGACTTGCTAACGGACT<br>AAAATCCTCGTGGATACGGC<br>CATGATAGCTATCTTCCGTC<br>CCGTTCATCTAAGGCAACAA<br>ACAGCTCAATTCCATCATCG<br>GAGTACGGCATTGATGAGTG<br>CCACAATGCCATGAAATTGC<br>AACTAAGCCAACGTCATCGC<br>CATCAATTTTCACATCGGCC<br>CCTATCGGATAGAACCCAAA<br>ATCCAAGTCAGACAGCAAGA<br>GGAGCCGTGTAAATTCGGAA<br>AGGCGTTATTCCTCAAATCA<br>TTGAAACGGTATTTGGCGGC<br>GGCAAAACGGCTTGTTTTGG<br>TTCGTATATAACGGTGGACC<br>TTTCTGTGGCAAGACAGGAC<br>CGGGCAAATAAGTGCGACTT<br>TGGAGACTACATTGCCTGAA<br>GGACCCACAGAATTTGTAGA<br>CCGTTGTGCTGGTTGAAAAT<br>GACCACTTGTCTCGTAATTT<br>GGGTTGATGGGTGGGAAATA<br>CGGGCATTCACATAAGTTTT<br>AACCTTGATGCTGATGATGC |

| | |
|---|---|
| *Krüppel* | CAGCTAATGCAGCAGCTAAG<br>CGCTTGTAGCTGATGTGTTG<br>CCATAGCTGGATAAATCGCA<br>ATTCCAAAAGCTGAGGCAGC<br>GGAAAGAGAGTGTTGGCCAA<br>GGCGAATGTAAATGCGTACC<br>TACCTAAAGGAGTGGACAGC<br>GTGCTGTTGGGGGAATTTAA<br>TGCTGATCTCGGTCTGAAAC<br>CCCGATGTATGGTACATATC<br>CACTGGAAGGCGGAGATATT<br>GTCGTGCGTTGAATTAGGAG<br>GACACATCCAGCATTTCCAC<br>GCAGATTTTACAGGTGAAGC<br>TGCTTATAGCCAAAGCTGCG<br>GACATTCGAAAGGCTTCTCA<br>AGTAAACCGCTTGTGGCACT<br>TTTCTCCAGTATGCAAACGC<br>GCAGTGCGAGCAATGATATG<br>CTCGCAAATGTCGTCTAAGA<br>TTTGCCATCGCAGATTTCAC<br>TTCGCACTCGAACGGCTTTT<br>CGTCGGAACTTCATGTGACA<br>CCACACTTGTGATTCATCAG<br>TGGATGCCTCGATAGCAATT<br>CACACATTGCCGCAAATCTA<br>ACTCATTCGAACCTCCGTAG<br>TGTCGCTTTTTCCATGTCGA<br>CTCCATCTTCAGACAAATCC<br>AAAACCCGACGAATGTCCTG<br>TTGCTCAGGCATATCACTGG<br>ATGCTCAAATCCTCTGGCTC<br>TGATGGCCCATATAAGAAGC |
| *knirps* | TAAGAGCGGCCAAAGAAGGA<br>CGCTGATGGTGCTGATGTTG<br>TTCTTCTTGTCGATGATGCA<br>CGTTGTAGCACTTCCTCAAG<br>TTGAACCAGTTGGAGCGACG<br>TGCTCCTGCAGCAGACAATG<br>AGCAGAGGCATATGTGGATG<br>CGGACAGATAGCTGGGATAG<br>CATCATGCTGAAGAAGGGCA<br>ATGGTAGCCTGGGAAGAGGA<br>TCAACGGAATCCACGCTCTG<br>TGCACGGAGTGAACATCCTC<br>CGACAAGCTCTGCATCTTGG<br>GCCAATGGAGCAAACCGAAA<br>GATGCACTGGTACAACGCTG |

| | |
|---|---|
| | CTTCTTGAGCGGAAACGGTG<br>TCTTCATGCTCAGATCCATG<br>TGTCGTTGAAGCTGTGCACG<br>TCCAGTTGGTAGAACTTCCG<br>ACACGAATATTCCCCTCATG<br>AACAACGAGGGTTTTTGGGG<br>GTGAGCGAGCACACGAATAA<br>TATCTCTTTCCACTTCCCTA<br>TTTCCAGCAAAACCTGTCTG<br>CAATGCAACTACCTGGTCAG<br>CCGCCGGCTAACTTTAAAAT<br>TGATTACTACTTCTACGGCT<br>ACAATAGCATTTGAGGGTGG<br>GCCAATCTTTAGACATACCA |

**Supplementary Table 2. Primer sequences.**

| Name | Sequences (5' to 3') |
|---|---|
| RT primer: Oligo(dT) | TTTTTTTTTTTTTTTTT |
| RT primer: P1-specific | GGCAAAGACCACATCCCG |
| RT primer: P2-specific | CTTACGAAAATCCCGACAAATTTGG |
| PCR forward primer: upstream of *hb* | TAAACATCGTGTCCAAACCCAT |
| PCR forward primer: P1-specific | CACTTGGACTGTCCGCAAGCCG |
| PCR forward primer: P2-specific | ACTGAGCGGCCACGAAACGGCC |
| PCR backward primer: upstream of *hb* | CTGCATTAACACTTCGTTCCCT |
| PCR backward primer: inside *hb*-RB | TTGGACTGTTGGTATTGTTTG |
| PCR backward primer: outside *hb*-RB | TCGTGGGCAAATATCTTAGACT |

**Supplementary Table 3. Sequences of HCR-FISH probes.**

| Target | Probe sequences (5' to 3') |
|---|---|
| *yellow*-B3 | GTCCCTGCCTCTATATCTCCACTCAACTTTAACCCGTACAAAAACTGCGGTCCATGTTTAT<br>GTCCCTGCCTCTATATCTCCACTCAACTTTAACCCGTACAACCACACTCATCCACTTTAAT<br>GTCCCTGCCTCTATATCTCCACTCAACTTTAACCCGTACAAGTATTTGGATTTGTGTCCAC<br>GTCCCTGCCTCTATATCTCCACTCAACTTTAACCCGTACAACCTCAATGGATCGGGGAAAA<br>GTCCCTGCCTCTATATCTCCACTCAACTTTAACCCGTACAAAAAATCCTCGTGGATACGGC<br>GTCCCTGCCTCTATATCTCCACTCAACTTTAACCCGTACAACCACAATGCCATGAAATTGC<br>GTCCCTGCCTCTATATCTCCACTCAACTTTAACCCGTACAAGGAGCCGTGTAAATTCGGAA<br>GTCCCTGCCTCTATATCTCCACTCAACTTTAACCCGTACAATTTCTGTGGCAAGACAGGAC<br>GTCCCTGCCTCTATATCTCCACTCAACTTTAACCCGTACAAGACCACTTGTCTCGTAATTT |
| *lacZ*-B5 | CTCACTCCCAATCTCTATCTACCCTACAAATCCAATAAAAATCACGACGTTGTAAAACGAC<br>CTCACTCCCAATCTCTATCTACCCTACAAATCCAATAAAAAAATGTGAGCGAGTAACAACC<br>CTCACTCCCAATCTCTATCTACCCTACAAATCCAATAAAAAGCTGATTTGTGTAGTCGGTT<br>CTCACTCCCAATCTCTATCTACCCTACAAATCCAATAAAAATTAACGCCTCGAATCAGCAA<br>CTCACTCCCAATCTCTATCTACCCTACAAATCCAATAAAAAATCGGTCAGACGATTCATTG<br>CTCACTCCCAATCTCTATCTACCCTACAAATCCAATAAAAATGCCAGTATTTAGCGAAACC<br>CTCACTCCCAATCTCTATCTACCCTACAAATCCAATAAAAATATTCGCTGGTCACTTCGAT<br>CTCACTCCCAATCTCTATCTACCCTACAAATCCAATAAAAACGGTTAAATTGCCAACGCTT<br>CTCACTCCCAATCTCTATCTACCCTACAAATCCAATAAAAAATCAATCCGGTAGGTTTTCC<br>CTCACTCCCAATCTCTATCTACCCTACAAATCCAATAAAAAATGTCTGACAATGGCAGATC |
| B3-H1-488 | CGGGTTAAAGTTGAGTGGAGATATAGAGGCAGGGACAAAGTCTAATCCGTCCCTGCCTCTA<br>TATCTCCACTC |
| B3-H2-488 | GTCCCTGCCTCTATATCTCCACTCAACTTTAACCCGGAGTGGAGATATAGAGGCAGGGACG<br>GATTAGACTTT |
| B5-H1-594 | ATTGGATTTGTAGGGTAGATAGAGATTGGGAGTGAGCACTTCATATCACTCACTCCCAATCT<br>CTATCTACCC |
| B5-H2-594 | CTCACTCCCAATCTCTATCTACCCTACAAATCCAATGGGTAGATAGAGATTGGGAGTGAGTG<br>ATATGAAGTG |

* For primary probes, black: initiator, blue: spacer, red: transcript specific sequences.
